# Selective Mitochondrial Respiratory Complex I Subunit Deficiency Causes Tumor Immunogenicity

**DOI:** 10.1101/2023.10.02.560316

**Authors:** Jiaxin Liang, Tevis Vitale, Xixi Zhang, Thomas D. Jackson, Deyang Yu, Mark Jedrychowski, Steve P. Gygi, Hans R. Widlund, Kai W. Wucherpfennig, Pere Puigserver

## Abstract

Targeting of specific metabolic pathways in tumor cells has the potential to sensitize them to immune-mediated attack. Here we provide evidence for a specific means of mitochondrial respiratory Complex I (CI) inhibition that improves tumor immunogenicity and sensitivity to immune checkpoint blockade (ICB). Targeted genetic deletion of the CI subunits *Ndufs4* and *Ndufs6*, but not other subunits, induces an immune-dependent tumor growth attenuation in mouse melanoma models. We show that deletion of *Ndufs4* induces expression of the transcription factor *Nlrc5* and genes in the MHC class I antigen presentation and processing pathway. This induction of MHC-related genes is driven by an accumulation of pyruvate dehydrogenase-dependent mitochondrial acetyl-CoA downstream of CI subunit deletion. This work provides a novel functional modality by which selective CI inhibition restricts tumor growth, suggesting that specific targeting of *Ndufs4*, or related CI subunits, increases T-cell mediated immunity and sensitivity to ICB.

## Introduction

Melanoma therapeutics have witnessed remarkable advancements over the past 15 years, transforming a previously untreatable disease into a more manageable one where some patients will be cured. Targeted combinatorial therapies such as BRAF and MEK inhibitors, have proven to be effective in patients harboring BRAF V600-missense mutations, although these cancers are generally prone to developing resistance to the therapy^1,2^. Additionally, immune checkpoint blockade (ICB) therapies such as anti-PD-1, anti-CTLA-4, and more recently anti-LAG3, have demonstrated remarkable efficacy in melanomas by blocking inhibitor signals that limit the ability of T-cells to kill cancer cells^3–7^. Despite these advances, many patients still do not meaningfully respond to ICB, highlighting the need to identify pathways and cell components that could be used to improve the immunogenicity of tumors^8^.

Targeting the mitochondria of tumor cells has been a promising area of therapeutic development over the last decade because many previous studies have implicated mitochondria in cancer cell survival, proliferation, and metastasis. Since mitochondria are a biosynthetic and bioenergetic hub inside of cells, many types of cancer cells, which proliferate quickly and have high energy demands, rely heavily on mitochondria for their survival^9,10^. In addition to survival, several studies have found that metastatic melanoma and breast tumors in a variety of animal models have increased oxidative phosphorylation (OxPhos) signatures^11–13^. OxPhos is driven by the electron transport chain, a series of five protein complexes made of over 100 individual proteins that drive ATP and electron equivalent synthesis in mitochondria^14^. Due to the prominent role of OxPhos and mitochondria in tumor growth and metastasis, significant efforts have been made to create OxPhos inhibitors, especially against CI, the NADH dehydrogenase complex. CI inhibitors have been very promising in the preclinical stage at treating a variety of cancer types^15–18^. However, once translated to the clinic, all inhibitors tested in humans have failed phase I clinical trials due to dose-limiting toxicities^19,20^. The success in preclinical models and significant animal and clinical genetic data pointing to a role for mitochondria in the survival and progression of cancer suggests that mitochondria and CI are promising targets; however, different approaches to inhibiting their activity need to be employed. We postulate that new modalities of selective targeting of CI could hold a larger therapeutic window in cancer treatment than current generation CI inhibitors.

In this work we adopted a more granular approach to selective targeting of CI subunits. Taking a genetic approach, we systematically knocked out many nuclear encoded subunits in the different domains of CI to examine its role in mediating an anti-tumor response in mouse melanoma models. We identify CI subunits, *Ndufs4 and Ndufs6*, that when deleted using CRISPR/Cas9 technology, induces an immune-dependent growth attenuation of melanoma tumors. Because previous attempts to inhibit CI focused on cell intrinsic growth inhibition, we prioritized understanding this novel cell extrinsic growth inhibition to introduce a new CI inhibition modality. Using this model, we show that selective CI inhibition through *Ndufs4* deletion can induce expression of MHC class-I antigen presentation and processing machinery, drive CD8^+^ and NK1.1^+^ cell responses to tumors and sensitize tumors to anti-PD-1 ICB. These studies suggest that selectively affecting CI structure, rather than broadly interfering with CI catalytic activity, can promote anti-tumor immunity, providing support for further research into mitochondrial control of tumor immunogenicity.

## Results

### Deletion of the CI subunits Ndufs4 and Ndufs6 causes an immune-dependent tumor growth attenuation

To study the effects of CI deficiency on tumor growth, we took a genetic approach and using CRISPR/Cas9 technology, knocked out several CI subunits from the three CI modules in B16-BL6 mouse melanoma cells. We then injected these cell lines into immune competent C57BL/6 and immunodeficient Fox1^nu^ (nude) mice, which lack functional T cells, to determine which genetic perturbations caused tumor growth attenuations and whether this attenuation was dependent on the adaptive immune system.

For all tested subunits in the Q-module or P-module of CI, responsible for transferring electrons to ubiquinone and pumping protons respectively^21^, deletion of these subunits had similar effects on tumor growth in both C57BL/6 and nude mice. In the Q-module sg*Ndufs3* cells had no tumor growth phenotype while sg*Ndufa9* cells exhibited mild tumor growth attenuation in both C57BL/6 and nude mice (Fig. 1a-c, Extended Data 1e). In the P-module sg*Ndufa1*, and sg*Ndufb8* exhibited similar growth defects in both C57BL/6 and nude mice (Fig.1d-f, Extended Data 1e). These data point to an immune independent mechanism of tumor growth attenuation that is likely caused by disrupted bioenergetics and or redox balance in these cells. This is further supported by blue-native polyacrylamide gel electrophoresis (BN-PAGE) analysis of these knockout cell lines that demonstrated a lack of CI activity and a lack of cell growth in galactose culture media (Extended Data 1a,d) and is in keeping with previous structural studies of CI^22^.

**Fig. 1:**
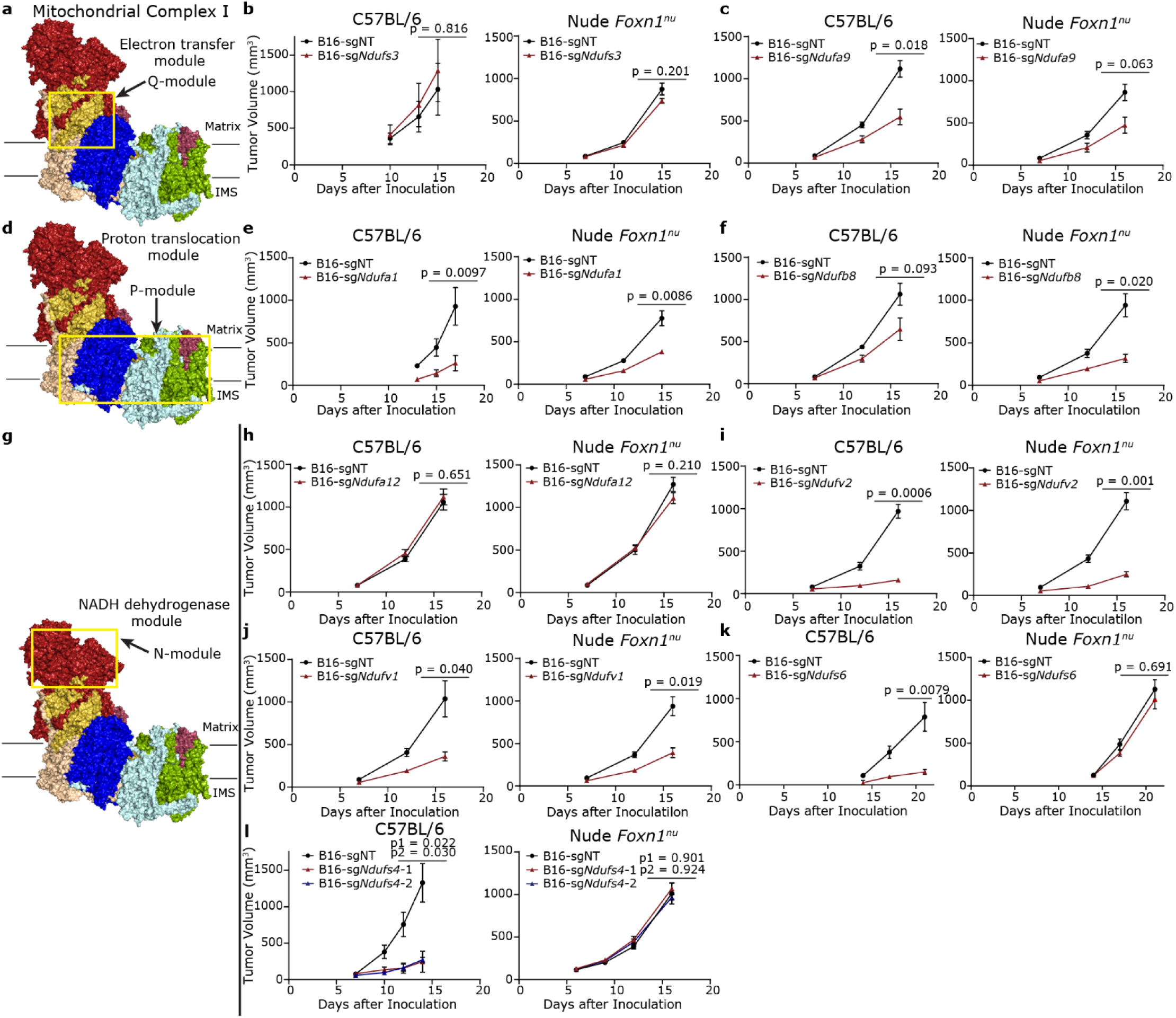
Deletion of mitochondrial CI components leads to cell intrinsic and cell extrinsic tumor growth attenuation in B16-BL6 melanoma tumors. **a**, Structure of murine CI with the Q-module highlighted (PDB: 6G2J), matrix refers to mitochondrial matrix while IMS refers to the inner membrane space^49^. **b**, Tumor growth curves of sgNT and sg*Ndufs3* tumors in C57BL/6 and nude mice. **c**, Tumor growth curves of sgNT and sg*Ndufa9* tumors in C57BL/6 and nude mice. **d**, Structure of murine CI with the P-module highlighted (PDB: 6G2J), matrix refers to mitochondrial matrix while IMS refers to the inner membrane space. **e**, Tumor growth curves of sgNT and sg*Ndufa1* tumors in C57BL/6 and nude mice. **f**, Tumor growth curves of sgNT and sg*Ndufb8* tumors in C57BL/6 and nude mice. **g**, Structure of murine CI with the N-module highlighted (PDB: 6G2J), matrix refers to mitochondrial matrix while IMS refers to the inner membrane space. **h**, Tumor growth curves of sgNT and sg*Ndufa12* tumors in C57BL/6 and nude mice. **i**, Tumor growth curves of sgNT and sg*Ndufv2* tumors in C57BL/6 and nude mice. **j**, Tumor growth curves of sgNT and sg*Ndufv1* tumors in C57BL/6 and nude mice. **k**, Tumor growth curves of sgNT and sg*Ndufs6* tumors in C57BL/6 and nude mice. **l**, Tumor growth curves of sgNT and sg*Ndufs4* tumors in C57BL/6 and nude mice. sg*Ndufs4*-1 and sg*Ndufs4*-2 denote different CRISPR guides against *Ndufs4*. P1 compares sgNT to sg*Ndufs4*-1 and P2 compares sgNT to sg*Ndufs4*-2. N= 5 mice per condition in **b**-**c**, **e**-**f**, and **h**-**i**. *P* values in **b**-**c**, **e**-**f**, and **h**-**k** were calculated using a Mann Whitney *u*-test, *p* values in **l** were calculated using the Brown-Forsythe and Welch ANOVA test with Dunnett’s T3 multiple comparisons test. Error bars are mean□±□s.e.m.

Most subunits in the catalytic NADH dehydrogenase N-module followed the same trend as the other modules. sg*Ndufa12* cells have no growth phenotype in C57BL/6 or nude mice, while sg*Ndufv1* and sg*Ndufv2* cells had attenuated tumor growth in both C57BL/6 and nude mice (Fig. 1g-j, Extended Data 1e). Similar to the other modules, there was a strong trend for subunits that caused growth defects to display a lack of CI activity and sensitivity to galactose (Extended Data 1a,d). However, deletion of the subunit *Ndufs4* with multiple CRISPR guides caused a stark tumor growth attenuation in C57BL/6 but not nude mice (Fig. 1l, Extended Data 1e), similar data was also observed knocking out *Ndufs6* (Fig. 1k). This unique phenotype indicated to us that there was a cell extrinsic, immune-dependent, tumor growth attenuation in these CI subunit deletion cell lines.

To better understand what was unique about deletion of *Ndufs4,* we assayed the assembly of CI and its activity. In keeping with previously published work^22^, knocking out *Ndufs4* resulted in altered CI assembly, but did not result in complete disassembly. As read out by NDUFA9, OXPHOS supercomplexes (co-assemblies of different stoichiometries of complex I, III and IV) in sg*Ndufs4* cells migrated slightly faster than those in sgNT or sg*Ndufa12* cells, which have previously been shown to not have an assembly defect^22^ (Extended Data Fig. 1b). Additionally, CI enzymatic activity could still be detected in sg*Ndufs4* cells using a BN-PAGE in-gel activity assay, albeit at a much lower molecular weight (Extended Data Fig. 1a,c). This indicates that the N-module is assembled in a much smaller complex, possibly by itself, and is partially functional. This partial functionality likely underlies the lack of growth defects in nude mice and the ability to grow in galactose culture media (Fig. 1l, Extended Data 1d).

### Ndufs4 deletion induces an anti-tumor response in multiple tumor models

To determine whether the anti-tumor immunity promoting effect of *Ndufs4* deletion was broadly applicable, we performed CRISPR-Cas9 mediated deletion of *Ndufs4* in another murine melanoma cell line, YUMM1.7 and the murine breast cancer line EO771. All cell lines displayed similar phenotypes to the B16-BL6 cell line, where deletion of *Ndufs4* leads to a large tumor growth attenuation in immunocompetent C57BL/6 mice but not immunodeficient nude mice (Fig. 2a-b). In addition to the marked attenuation of primary tumor growth, sg*Ndufs4* cells exhibited almost undetectable signs of melanoma lung metastasis. Strikingly, when injected into immunocompetent C57BL/6 mice via their tail vein, sg*Ndufs4* B16-BL6 cells did not form noticeable lung metastases and almost all the mice survived. However, in nude mice, tail vein injection of sgNT and sg*Ndufs4* B16-BL6 cells equally formed extensive lung melanoma metastasis and growth with similar survival rates (Fig. 2c,d).

**Fig. 2:**
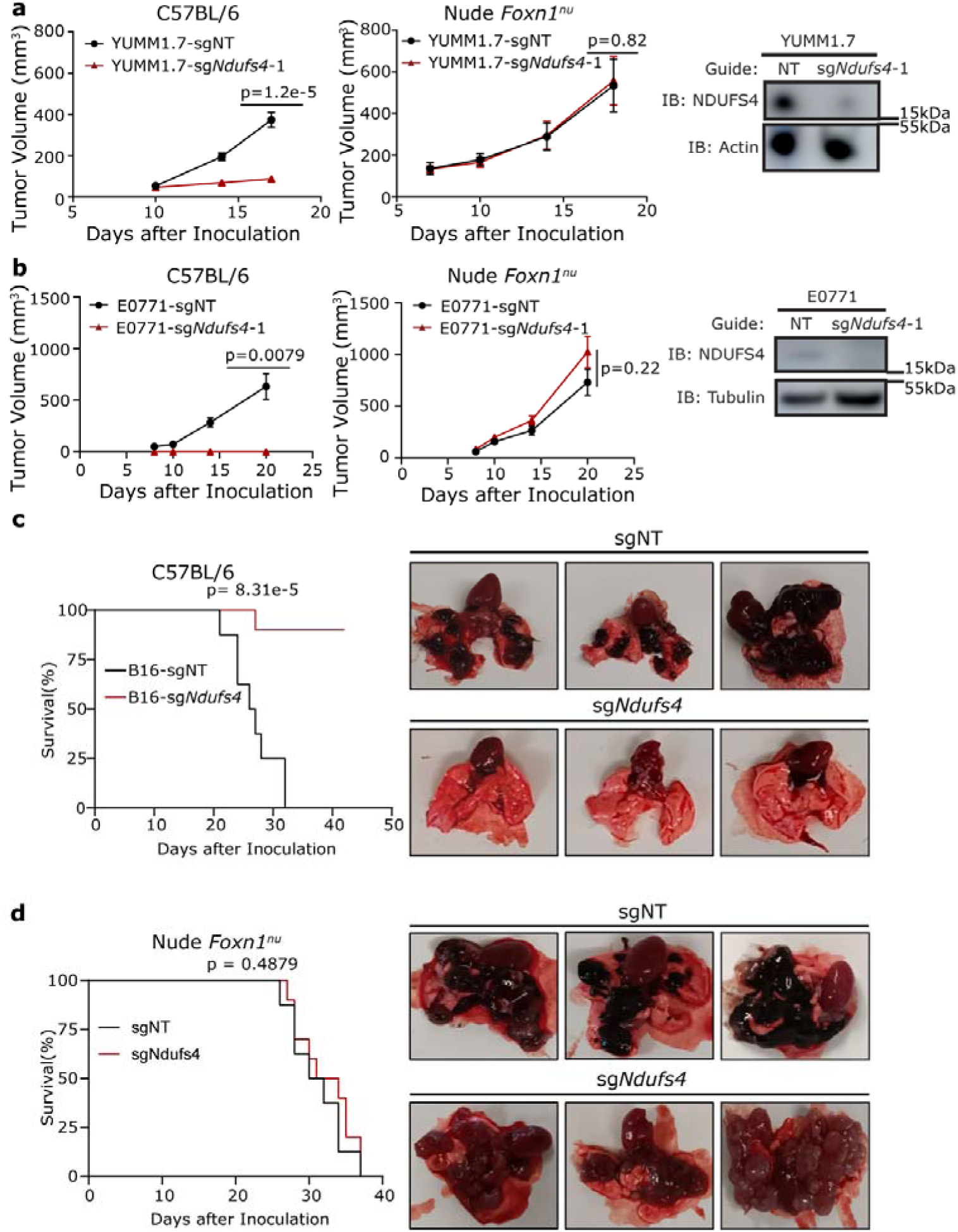
CRISPR/Cas9 mediated knockout of *Ndufs4* causes primary tumor growth defects in multiple cell line and attenuated metastasis in B16-BL6 cells. **a**, Tumor growth curves of YUMM1.7 sgNT and sg*Ndufs4* tumors in C57BL/6 and nude mice and western blot of sg*Ndufs4* expression. **b**, Tumor growth curves of E0771 sgNT and sg*Ndufs4* tumors in C57BL/6 and nude mice and western blot of sg*Ndufs4* expression. **c**, Kaplan-Meier plot of C57BL/6 mice injected via tail vein with sgNT or sg*Ndfus4* B16-BL6 cells. **d**, Kaplan-Meier plot of nude mice injected via tail vein with sgNT or sg*Ndufs4* B16-BL6 cells. N= 5 mice per condition in **a**-**b**, n=10 mice per condition in **c**-**d**. Representative pictures of lungs are shown in **c**-**d**. *P* values in **a**-**b** were calculated using a Mann Whitney *u*-test and *p* values in **c**-**d** using a Mantel-Cox log-rank test. Error bars are mean□±□s.e.m.

To ensure that the effects on tumor growth we were seeing were due to the loss of *Ndufs4* and not an off-target effect of the CRISPR guides, we overexpressed *Ndufs4* or a control plasmid (V300) in sgNT and sg*Ndufs4* B16-BL6 cells. We observed a marked rescue in growth in the sg*Ndufs4* cells re-expressing *Ndufs4,* while the over expression of *Ndufs4* had no effect on the tumor growth of sgNT cells (Extended Data Fig. 2a). While the rescue of tumor growth was not complete, expression of *Ndufs4* in the re-expression cell line was still lower than in sgNT cells and there isn’t a statistically significant difference between the tumor growth of the sgNT and the *Ndufs4* re-expressing cells.

Mutations in CI in humans and mouse models have been shown to cause inflammation and cell death^23,24^. Prior work has shown that some of this inflammation and cell death can be attributed to altered NAD/NADH ratios resulting from CI deficiencies^23^. To test whether these immunogenic effects could be caused by a change in redox state in the cell we overexpressed a NADH oxidase from *Lactobacillus brevis*, LbNOX, which we have previously shown to rescue cell death in mitochondrial disease models^23^. Overexpression of the cytosolic or mitochondrial targeted versions of the LbNOX protein did not alter the growth of sgNT or sg*Ndufs4* tumors (Extended Data Fig. 2b). Therefore, we conclude that the immunogenicity that is generated by the sg*Ndufs4* cells is not a consequence of NADH-dependent redox imbalance, rather another role that CI is playing in the tumor cells.

### sgNdufs4 tumors show greatly increased antigen processing and MHC class-I proteins in a cancer cell intrinsic manner

To investigate the underpinning of the immunogenicity of sg*Ndufs4* cells, we performed quantitative TMT-proteomics on tumors from C57BL/6 and nude mice, identifying almost 8,000 proteins (Fig. 3a, Extended Data Fig. 3a-d). In the sg*Ndufs4* tumors grown in C57BL/6 mice, we observed a stark enrichment of immune related gene ontology (GO) processes among the upregulated proteins (Extended Data Fig 3a). While this enrichment isn’t observed in the nude mice (Extended Data Fig. 3b), many antigen presentation process proteins are similarly upregulated in sg*Ndufs4* tumors from the nude mice (Fig. 3b, red asterisk) including NLRC5, H2-K1, PSMB8, PSMB9, PSMB10 and TAP1. These upregulated proteins span all steps in the antigen presentation process. We observe increased levels of the MHC class I proteins needed to present peptide antigens (H2K1), the peptide transporters responsible for carrying protein peptides to the endoplasmic reticulum where they are loaded onto MHC class I molecules (TAP1), the immunoproteasome subunits (PSMB8-10) which efficiently generate MHC class I ligands and the transcription factor NLRC5 that has been shown to regulate all these genes^25^. The upregulation of these proteins across the entire antigen presentation pathway and previous literature points to NLRC5 as a potential mediator of the observed increase in antigen presentation machinery. Using qPCR, we were able to validate that many of the genes encoding the enriched proteins in the proteomics were increased at the mRNA level in both B16-BL6 and YUMM1.7 tumors (Extended Data Fig. 4a,b). Because these proteins are upregulated in both nude and C57BL/6 mice, it suggests that this increase is primarily driven by cell intrinsic mechanisms, rather than simply reflecting increased immune cell invasion or activation. Furthermore, by qPCR, the expression of *Nlrc5* and its target genes *H2-K1, Psmb8, Psmb9, Psmb10* and *Tap1* are upregulated in sg*Ndufs4* cells grown in culture (Fig. 3c), further supporting the hypothesis that these increases in antigen processing and presentation are cancer cell intrinsic.

**Fig. 3:**
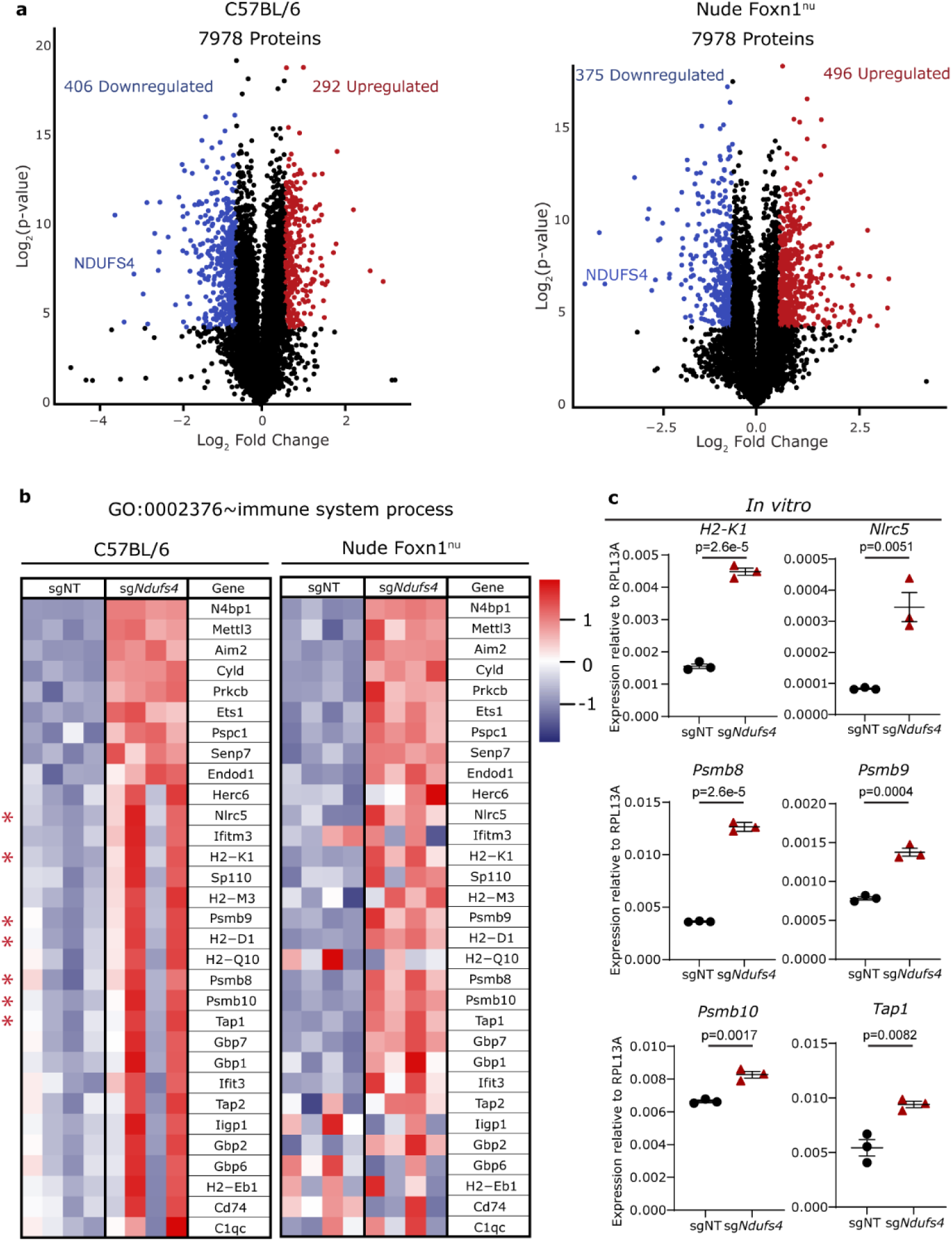
MHC-I and antigen processing proteins are increased in sg*Ndufs4* tumors and cells in a cell intrinsic manner. **a**, Volcano plots of proteomics done in sgNT and sg*Ndufs4* tumors in C57BL/6 and nude mice (n=4 tumors per condition). NDUFS4 is labeled as a positive control. **b**, Heatmap of Gene Ontology term 0002376 in proteomic dataset of sgNT and sg*Ndufs4* tumors in C57BL/6 and nude mice. Asterisk denotes MHC-I and antigen processing genes. **c**, qPCR mRNA levels of MHC-I and antigen processing genes of sgNT and sg*Ndufs4* cells cultured *in vitro*. *P* values in **c** were calculated using a two-tailed, unpaired Student’s t-test. Error bars are mean□±□s.e.m.

### sgNdufs4 tumor immune signatures correlate with better clinical outcomes

To determine if the immune signatures of increased antigen processing and presentation we observed in sg*Ndufs4* tumors correlate with clinical outcomes, we surveyed the Skin Cutaneous Melanoma (SKCM) TCGA and melanoma immune therapy response datasets. In the SKCM dataset, higher expression of the human homologs of individual genes found to be significantly upregulated in sg*Ndufs4* tumors such as *NLRC5, HLA-A* (*H2-K1* homolog), *PSMB8, PSMB9,* and *PSMB10* all correlate with longer patient survival (Fig. 4a). Additionally, RNA expression of *NLRC5, HLA-A* (*H2-K1* homolog), *TAP1, PSMB8, PSMB9,* and *PSMB10* all tended to be higher in metastatic melanoma patients who responded to ICB versus non-responders (Fig. 4b). These data give a strong indication that the immune signature induced in sg*Ndufs4* tumor cells is clinically relevant in multiple contexts.

**Fig. 4:**
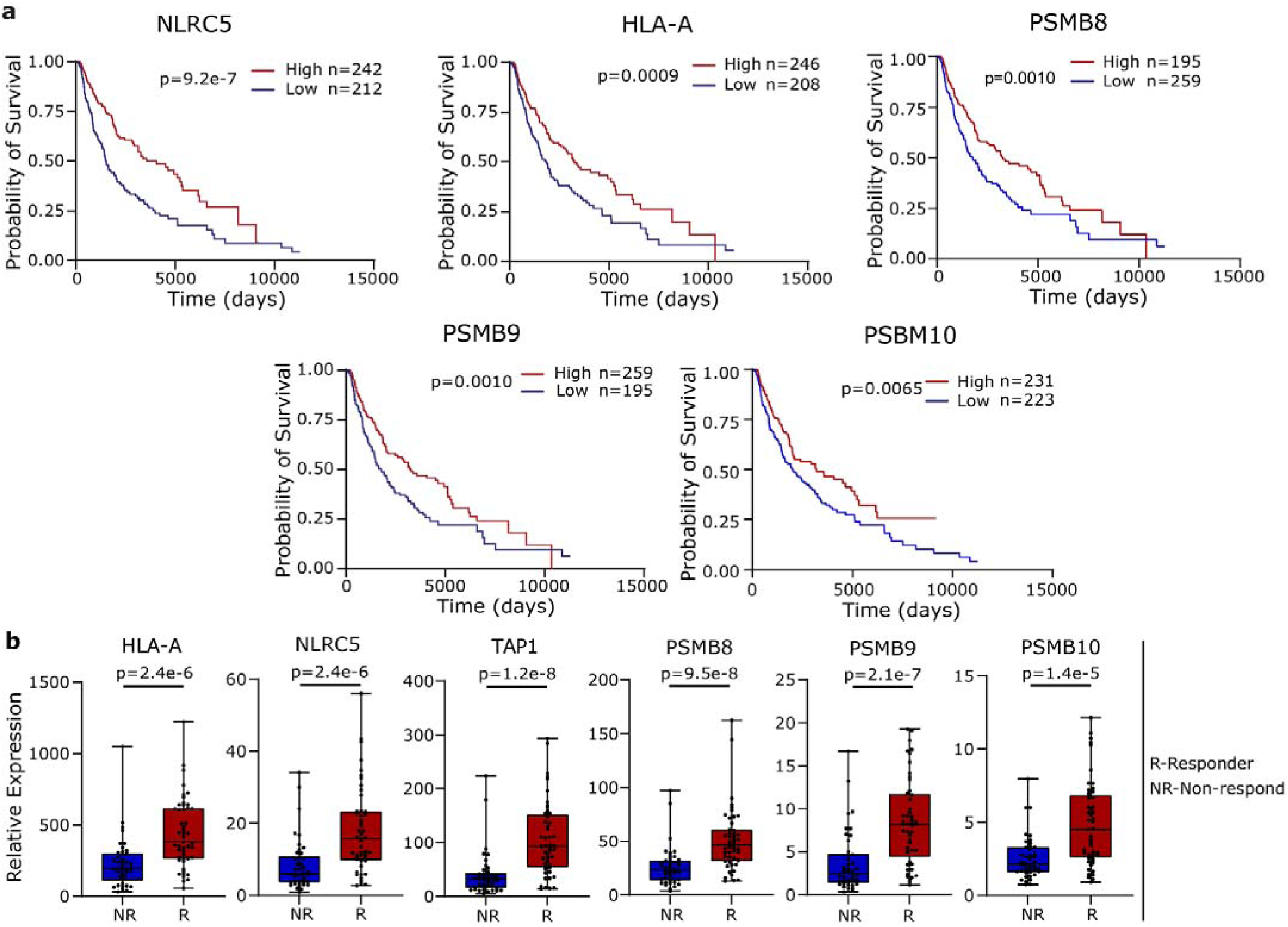
The cell intrinsic immune signature of sg*Ndufs4* cells correlates with improved prognosis and responsiveness to immunotherapy in melanoma patients. **a**, Kaplan-Meier curves of SKCM TCGA patients segregated by the top and bottom 50% of expression of human MHC-I and antigen processing machinery homologs that are upregulated in sg*Ndufs4* cells. **b**, RNA expression values of human MHC-I and antigen processing machinery homologs that are upregulated in sg*Ndufs4* cells in melanoma patients segregated by patients response to immunotherapy (data from PMID: 30753825^50^, accessed through TIGER^48^). *P* values in **a** were calculated using a Mantel-Cox log-rank test and *P* values in **b** were calculated using a Mann Whitney *u*-test. Box and whisker bars represent min to max with the box representing the interquartile range with the median plotted.

Additionally, while SKCM displayed the strongest correlation with this sg*Ndufs4* immune signature (Extended Data Fig. 5), several other cancer types also strongly correlate with the antigen processing and presentation genes, including bladder urothelial carcinoma, mesothelioma, sarcoma, ovarian serous cystadenocarcinoma, and breast invasive carcinoma (Extended Data Fig. 5), suggesting that these mechanisms of immune activation could be applicable in other cancer types.

### Nlrc5 and target gene expression is driven by altered mitochondrial metabolism

To better understand the mechanism by which *Ndufs4* deletion drives increased *Nlrc5* gene expression, we first tested several other pathways that had previously been shown to induce *Nlrc5*. First, because *Nlrc5* expression was originally found to be driven by promoter DNA methylation^25^, we performed bisulfate sequencing on the *Nlrc5* promoter region in sgNT and sg*Ndufs4* cells in vitro. We found that both sgNT and sg*Ndufs4* cells had heavily methylated *Nlrc5* promoters, but there were no significant methylation differences between the two cell types (Extended Data Figure 6a). Second, given that NF-κB activation has been shown to induce *Nlrc5* expression^26^, we tested the dependency of our phenotype on several targets of NF-κB. A primary activator of NF-κB, especially in the case of mitochondrial disruption, is cGAS-STING, a cytosolic DNA sensor. Under basal conditions, a downstream phosphorylation target of cGAS-STING, TBK1, is not appreciably phosphorylated in either sgNT or sg*Ndufs4* cells (Extended Data Figure 6b). Additionally, neither STING nor a downstream transcription factor STAT1 are necessary for tumor immunogenicity (Extended Data Figure 6c,d). Lastly, inflammasome activation is often downstream of NF-κB activation and the inflammasome component Aim2 was increased in our proteomics, we also tested whether inflammasome activation is detectable using IL-1β as a readout. In ELISAs performed on cell culture media, no IL-1β was detectable in either sgNT or sg*Ndfus4* cells, indicating that inflammasome activation likely doesn’t mediate the immune phenotype (Extended Data Figure 6e).

We next searched for alternative mechanisms through which altered mitochondrial activity could modulate *Nlrc5* transcription, as it appeared that previously described mechanisms were not relevant to our system. Because many mitochondrial derived metabolites such as NADH/NAD+, acetyl-CoA, succinate and fumarate have previously been implicated in retrograde signaling from the mitochondria to the nucleus^27–29^, we performed mitochondrial metabolomics on sgNT and sg*Ndufs4* melanoma cells. These experiments showed that sg*Ndfus4* cells contained increased levels of acetyl-CoA and decreased levels of the TCA cycle intermediates aconitate, malate, oxaloacetate, succinate, and succinyl-CoA (Fig. 5a-c). This suggests that deletion of *Ndufs4* is inhibiting flux through the TCA cycle, in line with previous work demonstrating that inhibition of CI activity causes reduced TCA flux^27,30^. To test whether this increase in mitochondrial acetyl-CoA was the cause of gene expression changes, we inhibited acetyl-CoA formation from pyruvate using Devimistat, a pyruvate dehydrogenase (PDH) inhibitor and performed qPCR analysis on *Nlrc5* and its target genes. Inhibition of PDH *in vitro* dramatically reduced the expression of *Nlrc5* and its target genes in both B16-BL6 and EO771 sg*Ndfus4* cells but had little effect on sgNT cells (Fig. 5d-e). These data suggest that accumulation of acetyl-CoA in the mitochondria is leading to these gene expression changes we observe in sg*Ndufs4* cells. Because mitochondrial acetyl-CoA is the primary substrate for histone acetylation^31^, it is conceivable that deletion of sg*Ndufs4* mediates *Nlrc5* gene expression likely through acetyl-CoA dependent epigenetic reprogramming. However, the exact histone acetyltransferases or other proteins involved in this remodeling and the extent of it remains unknown.

**Fig. 5:**
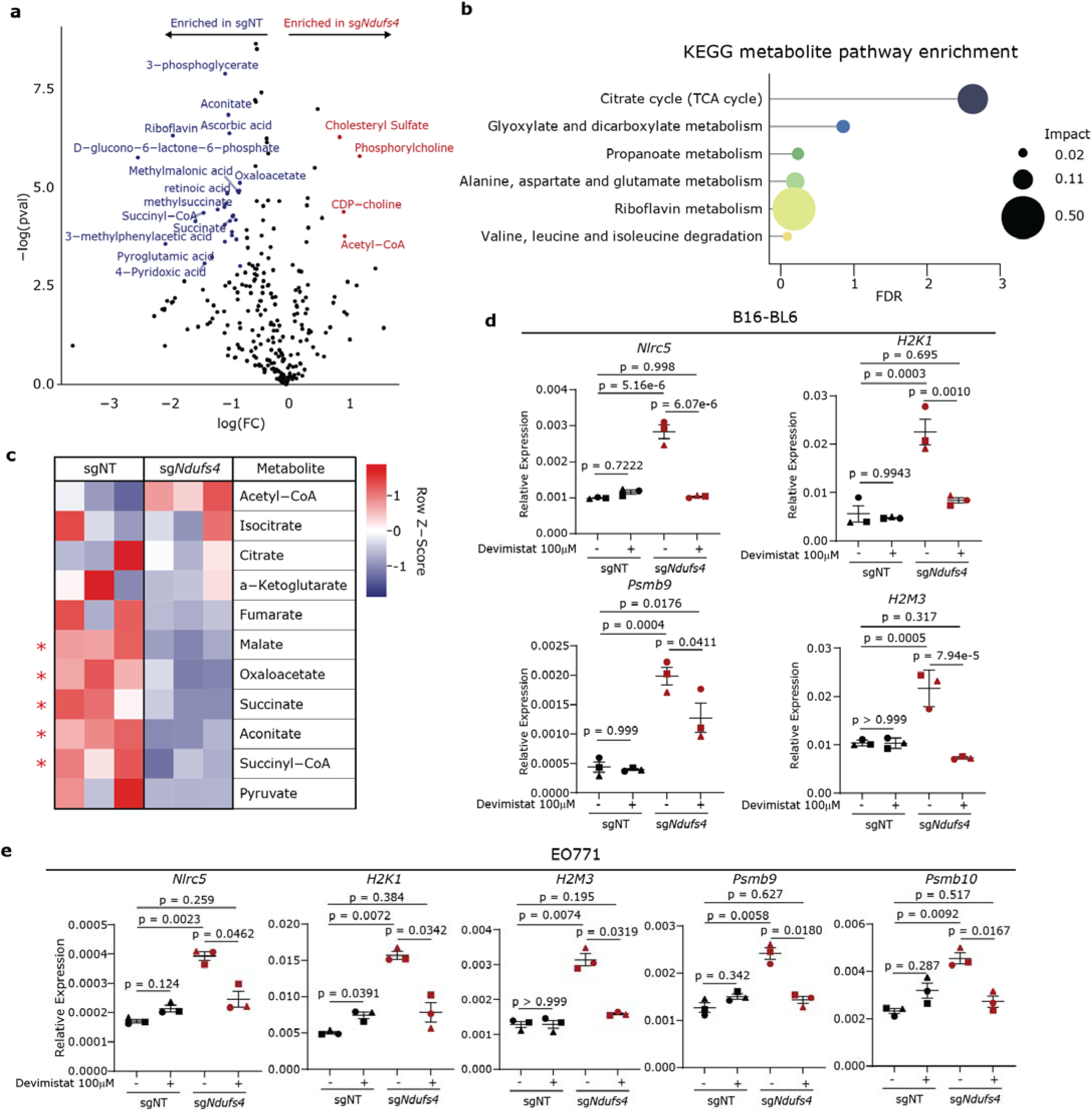
sg*Ndufs4* cells have decreased TCA cycle intermediates and increased acetyl-CoA which drives *Nlrc5* gene expression. **a**, Metabolomics from crude mitochondria isolated from sgNT and sg*Ndufs4* cells in culture. **b**, KEGG metabolite pathway enrichment analysis of metabolites with fold-change of >0.5 in sg*Ndufs4* cells and a *p*-value<0.05. **c**, Heatmap of all TCA cycle metabolites in sgNT and sg*Ndufs4* cells, red asterisk denotes *p*-value<0.05 on unpaired Student’s *t*-test. **d**, qPCR of *Nlrc, H2-K1, PSMB10 and H2M3* in sgNT and sg*Ndufs4* B16-BL6 cells treated with DMSO or 100μM Devimistat (PDH inhibitor). **e**, qPCR of *Nlrc, H2-K1, PSMB9, PSMB10 and H2M3* in sgNT and sg*Ndufs4* EO771 cells treated with DMSO or 100μM Devimistat (PDH inhibitor). *P*-values in **a** were calculated using a two-tailed Student’s *t*-test. For **d**-**e** shapes indicate averages of independent replicates, *p*-values are from Brown-Forsythe and Welch ANOVA test with Dunnett’s T3 multiple comparisons test.

### sgNdufs4 tumors are enriched with activated CD8^+^ T-cells and NK cells and respond better to anti-PD-1 therapy

In order to understand the nature of the immune response mounted against sg*Ndufs4* tumors, we performed fluorescence-activated cell sorting (FACS) analysis on the CD45^+^ cells in the tumor microenvironment (TME). In the T-cell panel, there was no change in the total number of CD8 or CD4 T-cells in the TME (Fig. 6a, Extended Data Fig. 7a). However, there were significantly more CD44^+^CD69^+^ double positive and PD-1^+^TIM3^+^ double positive CD8^+^ T-cells, indicating that the T-cells in the TME were significantly more activated (Fig. 6a, Extended Data Fig. 7a). This result is consistent with our proteomics showing an increase in the MHC I allele protein levels in the sg*Ndufs4* tumors.

**Fig. 6:**
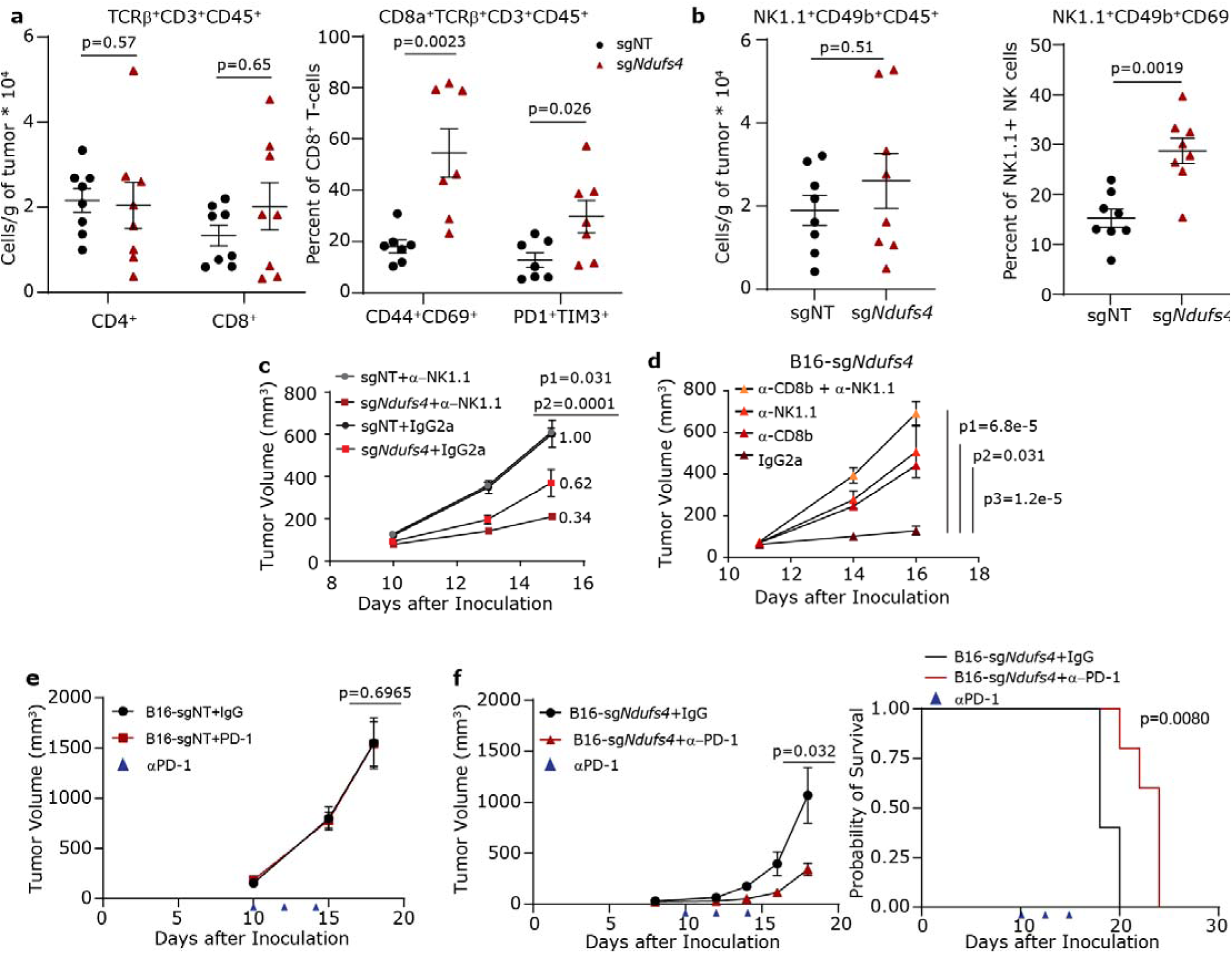
The immune response against sg*Ndufs4* tumors is dependent on both CD8b^+^ and NK1.1^+^ cells and synergizes with PD-1 therapy. **a**, Quantification of FACS based analysis of CD4^+^ and CD8^+^ T-cell infiltration and CD8^+^ T-cell activation in sgNT and sg*Ndufs4* tumors grown in C57BL/6 mice. N = 8 mice per condition, representative plots shown of two independent experiments. **b**, Quantification of FACS based analysis of NK cell infiltration and activation in sgNT and sg*Ndufs4* tumors grown in C57BL/6 mice. n = 8 mice per condition, representative plots shown of two independent experiments. **c**, Tumor growth curves of sgNT or sg*Ndufs4* tumor grown in C57BL/6 mice grown treated with either control IgG or anti-NK1.1 antibodies. P1 compares sgNT+anti-NK1.1 to sg*Ndufs4*+anti-NK1.1 and p2 compares sgNT+IgG to sg*Ndufs4*+ IgG. **d**, Tumor growth curves of sg*Ndufs4* tumors grown in C57BL/6 mice treated either with control IgG, anti-CD8b, anti-NK1.1 or both anti-CD8b and anti-NK1.1 antibodies. P1 compares sg*Ndufs4+*IgG to anti-CD8b+NK1.1, p2 compares sg*Ndufs4*+IgG to anti-NK1.1, p3 compares sg*Ndufs4*+IgG to anti-CD8b. **e**, Tumor growth curves of sgNT tumors in C57BL/6 mice treated either with a control IgG or anti-PD-1. **f**, Tumor growth curves and corresponding survival plot of C57BL/6 mice bearing sg*Ndufs4* tumors and treated either with a control IgG or anti-PD-1 antibodies. *P* values in **a**, **b**, **e**, **f**, were calculated using a Mann Whitney *u*-test, *p* values in **c** were calculated using two-way ANOVA followed by Tukey’s post-hoc test, while *p* values in **d** were calculated using one-way ANOVA followed by Tukey’s post-hoc test. Error bars are mean□±□s.e.m.

Similar to the T cell phenotype, we found that there was no significant difference in the number of NK cells in the sg*Ndufs4* TME (Fig. 6b, Extended Data Fig. 7b). However, tumor resident NK cells display greatly increased levels of the activation marker CD69 (Fig. 6b, Extended Data Fig. 7b). Additionally, by qPCR, we see a significant increase in the mRNA levels of immune related genes such as *Ifng*, *Prf1,* and *Gzmb* in sg*Ndufs4* B16-BL6 and YUMM1.7 tumors from C57BL/6 mice (Extended Data Fig. 7c,d), further supporting a model of immune activation in the sg*Ndufs4* TME. Together these data support a model of an inflamed TME that is leading to the enhanced activation of different immune cell populations and increased killing of sg*Ndufs4* tumor cells.

To test whether NK1.1^+^ and CD8b^+^ cells were required for the sg*Ndufs4* immunogenicity, we immunodepleted NK1.1^+^ and CD8β^+^ cell types using anti-NK1.1 and anti-CD8β antibodies, respectively. In the individual antibody treatments, we saw similar and marked rescues of tumor growth for both NK1.1 and CD8b antibodies (Fig. 6c,d). The combination treatment of NK1.1 and CD8b was more effective than either antibody in isolation, again indicating that CD8β^+^ and NK1.1^+^ cells independently contribute to the observed immunogenicity.

Because of the increased T cell activation and MHC-I expression in sg*Ndufs4* tumors, we hypothesized that sg*Ndufs4* tumors would be sensitive to immunotherapies. sgNT B16-BL6 tumors showed no response to anti-PD-1 therapy (Fig. 6e), in line with previous work showing that mice with B16-BL6 tumors only meaningfully respond to immunotherapy after being vaccinated with either the GM-CSF or Flt3-ligand^32^. However, mice bearing sg*Ndufs4* tumors lived longer and had substantially smaller tumors on average when treated with anti-PD-1 therapy (Fig. 6f). Taken together these results show that a modality of selective inhibition of CI leads to an immune-mediated anti-tumor response and renders mouse melanoma models more sensitive to immunotherapies.

## Discussion

Together, this work has identified a novel modality of mitochondrial CI inhibition in tumors that leads to an immune-dependent attenuation of tumor growth. This mode of CI inhibition, which is specifically triggered by deletion of the CI subunit *Ndufs4,* increases the activation status of CD8^+^ and NK cells within the tumor environment and synergizes with anti-PD-1 ICB. This finding has potential implications in the field of immunotherapy. While many cancer patients, especially with metastatic melanoma, respond to current generation ICB therapies, a large proportion of patients still have a limited response to the treatment^33^. Therefore, any therapies that could boost the effectiveness of current ICB therapies should be investigated for the benefit of future patients. Additionally, because the highlighted mechanism of action of CI inhibitors as cancer therapeutics has previously focused on cell intrinsic apoptosis^34^, this novel link between selective CI inhibition and induction of tumor immunogenicity opens up a new potential class of co-therapy development.

The discovery of new mechanisms to regulate MHC class-I expression in tumors is also of great interest to the field of immunotherapy. In order for ICB to be successful, patients need to be able to develop and sustain a T-cell based immune response against their cancer prior to ICB, as ICB can only stimulate a preexisting T-cell response^35^. Therefore, the downregulation of MHC-I molecules by tumor cells to evade T-cell surveillance has been linked to both intrinsic and acquired ICB resistance in multiple systems^36,37^. Thus, the development of therapies that can induce MHC-I molecules and antigen processing machinery is an active area of research. We present a novel mechanism by which selective partial CI inhibition can induce MHC-I antigen presentation and processing machinery through mitochondrial nuclear metabolic signaling.

Mitochondria have long been seen as a metabolic signaling hub in cells for bioenergetics, redox status, metabolism and inflammation^38–41^. Here we show that alterations in mitochondrial TCA-related metabolism can lead to changes in immunogenic-dependent tumor growth inhibition. However, unlike most previously published examples such as succinate, fumarate, aconitate and alpha-ketoglutarate^42^, the effect of the metabolite changes seems to be centered on the tumor cells versus immune cells. While mitochondria have been widely implicated in inflammatory signaling, less work has demonstrated the ability of mitochondrial metabolites to induce a pro-inflammatory tumor environment. Previous works have focused much more on mitochondria as a signaling platform for MAVS/RIG-1 signaling for RNA recognition, mtDNA recognition by cGAS-STING leading to NF-κB activation, or how mitochondrial dysfunction can activate the inflammasome^39^. However, a recent publication has shown that inhibition of mitochondrial CII can promote MHC class-I antigen processing transcription through a succinate driven epigenetic mechanism^43^. However, they claimed that CI had no such role in regulating antigen presentation pathways. Here we show that selective CI inhibition can also promote similar transcriptional changes through a distinct mechanism centered upon accumulation of acetyl-CoA. To our knowledge these are the first two examples of how altering mitochondrial electron transfer chain metabolism can drive anti-tumor immunity in a cell intrinsic manner in tumor cells. Taken together these works establish mitochondrial metabolism as a new potential therapeutic target in cancers with low MHC class-I expression.

## Acknowledgements

Claudia Adams Barr program in Cancer Research (to P.P.), Dana-Farber Cancer Institute Innovation Research Funds (to P.P.), NIH R01CA181217 (to P.P.), DOD CDMRP W81XWH-22-1-0780 (to P.P.), Melanoma Research Foundation (to PP), CRI Fellowship CRI4166 (to J.L.), R01 CA238039, R01 CA251599, R01 CA234018, P01 CA163222 and P01 CA236749 (to K.W.W.), the Ludwig Center at Harvard Medical School (to K.W.W.) and an AACR-Merck Immunooncology Research Fellowship 22-40-68-YU (to D.Y), Beyond the Sun Drenched Skies philanthropic fund (to H.R.W.). K.W.W. is co-director of the Parker Institute for Cancer Immunotherapy (PICI) at Dana-Farber Cancer Institute.

## Disclosures of Conflict of Interest

K.W.W. serves on the scientific advisory boards of TScan Therapeutics, Bisou Bioscience Company, DEM BioPharma, Solu Therapeutics and Nextechinvest, and he receives sponsored research funding from Novartis. K.W.W. is a co-founder, stockholder and advisory board member of Immunitas Therapeutics, a biotech company.

## Methods

### Animals and cancer cell lines

Homozygous outbred nude mice (Jackson Lab: Foxn1 nu /Foxn1 nu #007850) and C57BL/6 (Jackson Lab: #000664) were used for xenograft tumor experiments under the auspice of Beth Israel Deaconess Medical Center animal facility and IACUC approved protocols. YUMM1.7 was a kind gift from Marcus Bosenberg (Yale University School of Medicine). B16-BL6 was a kind gift from James Allison (MD Anderson Cancer Center). All cell lines were maintained in a humidified incubator at 37°C with 5% CO2, and if not otherwise indicated, in DMEM (Sigma-Aldrich) with 10% FBS, 100 U/ml penicillin, and 100 mg/ml streptomycin.

### Western blot assays

For Western immunoblotting, cells were lysed in RIPA buffer at indicated time points, and the protein concentration was quantified using the DC protein concentration assay (Pierce) before being subjected to gradient 4-12% (30:1) SDS polyacrylamide electrophoresis and subsequently transferred to PVDF membranes. Specifically, MES SDS running buffer was used in gels to detect proteins whose molecular weight are below 20 kD, while MOPS SDS running buffer was used for all other Western blot assays. Membranes were blocked with 2% BSA in TBST, then probed with primary antibodies overnight, and subsequently using secondary antibodies for 1 h at room temperature.

### Quantitative real-time PCR (qPCR)

RNA was isolated with Trizol (Invitrogen, 15596-026) and the Zymo-Spin Direct-zol RNA Kit (Zymo Research, R2050). 1 μg of RNA was used to generate cDNA with a High-Capacity cDNA Reverse Transcription Kit (Applied Biosystems, 4368813) following the manufacturer’s protocol. For gene expression analysis, cDNA samples were mixed with Sybr Green quantitative PCR master mix (Applied Biosystems, 4309155) and were analyzed by a CFX 384 Real-Time system (Bio-Rad).

### Plasmid construction, lentiviral generation, and transduction

The plasmid for overexpression of Ndufs4 was purchased from VectorBuilder with the guide RNA targeted sequence synonymously altered. CRISPR/Cas9 mediated gene knockout was performed using the GeCKO system, where pLentiCRISPRv2 (Addgene, 98290) was digested with BbsI enzyme and pre-annealed 5’-end phosphorylated sgDNA sequences (found in Extended file 2) inserted using Quick Ligase (New England Biolabs), and subsequently transformed into Stabl3 TM E. coli. Resulting plasmid inserts were verified by sequencing (Genewiz). Replication-defective lentiviral supernatants were generated using transfection of 600ng psPAX2 (Addgene, #12260), 300ng pMD2.G (Addgene, #12259) and 900ng lentiviral plasmid backbone using PolyFect (Qiagen, 301105) into HEK293t cells in 6-well format according to the manufacturer’s instructions. Supernatants collected twice (at 48 and 72h post-transfection, and filtered through a 0.45-µm filter, then used to transduce cells in the presence of 8 μg/ml polybrene. The transduced cells were then selected with 2 μg/ml of puromycin or 8 μg/ml blasticidin for 4 days and then cultured without antibiotics for at least 7 days prior to use in experiments.

### Tumor allograft studies using melanoma and breast cancer cell lines

Tumor cells (1×10 5 cells for B16-BL6 and Yumm1.7 melanoma and EO7071 breast cancer cells) were injected subcutaneously into the flanks of 7-week-ld female C57BL/6 mice or nude mice with a 30G needle. The tumor volumes were measured by a caliper and calculated as (Volume = length x width 2 / 2). The experiments were when the tumor volumes reach around 2000 mm 3 or any sign of suffering of mouse, such as decreased activity, 20% body weight loss, back-arching, etc. The tumor samples at the final time points were collected for proteomics analyses and infiltrated immune cells analyses by flow cytometry. For the depletion of NK1.1 + and CD8 + cells in mouse, InVivoMAb anti-mouse NK1.1 (Bioxcell, Catalog# BE0036, Clone: PK136) and InVivoMAb anti-mouse CD8β (Lyt 3.2) (Bioxcell, Catalog# BE0223 Clone: 53-5.8) were administrated at 0.1 mg per mice on one day before melanoma cell subcutaneous inoculation and then twice per week. The tumor volumes were closely monitored. Spleens were collected for the analysis of NK1.1 + and CD8 + cells to validate the efficacy of antibody treatment.

### In vivo metastasis assay

50 thousand B16-BL6 cells with/without Ndufs4 deletion in cold DMEM medium were tail-vein injected into 7-week-old female C57BL/6 mice or nude mice. The mice were closely monitored for their health. The mice were sacrificed if any signs of suffering of mouse were observed, such as decreased activity, 20% body weight loss, back-arching and short-breathing. The sacrificed mice were dissected to check and photograph lung metastasis status. The last days of mice were recorded and used for the analysis of survivals.

### Flow cytometry

Tumors were harvested on day 16 following inoculation, weighed, and dissected into small pieces using sterile scalpels in serum-free RPMI 1640 media. Tumor fragments were dissociated in 160 U/mL Collagenase Type IV (Gibco), 80 U/mL DNase I (Sigma), 0.1 mg/ml Hyaluronidase Type V (Sigma) using GentleMACS M tubes on the GentleMACS™ Dissociator (Miltenyi Biotec #130-093-235) followed by incubation at 37°C for 30 min. Following enzymatic digestion, the reaction was quenched with FACS buffer and passed through 70µm cell strainers, and then centrifuged at 300 × *g* for 6 minutes to pellet the cells. CD45+ cells were enriched using CD45 TIL microbeads (Miltenyi Biotec) following the manufacturer’s protocol. For flow cytometry analysis, cells were stained for 15 minutes at RT with 100 μL of Zombie UV dye (BioLegend) diluted 1:500 in PBS. Cells were washed once with FACS buffer and pre-incubated with 5 μg/mL TruStain FcX anti-mouse CD16/32 antibody (clone 93, BioLegend) in 100 μL FACS buffer for 5 minutes on ice before immunostaining. For flow cytometry analysis of T cells, NK cells and NKT cells, cells were incubated with a combination of fluorochrome-conjugated antibodies to the following surface markers (from Biolegend unless otherwise indicated): CD3ε (17A2, Biolegend/ BD Bioscience), TCRβ (H57-597, Biolegend/ BD Bioscience), CD4 (RM4-4), CD8α (53-6.7), CD44 (1M7), CD45 (30-F11), CD62L (MEL14), CD69 (H1.2F3), NK1.1 (PK13), CD49b (HMα2), NKG2D (CX5), PD1 (29F.1A12), Tim3 (RMT3-23). The cells were stained in 100µl FACS buffer for 15 minutes at RT, washed twice with FACS buffer and subsequently fixed in either Fixation Buffer (BD Biosciences) or fixed and permeabilized using the FoxP3/Transcription Factor Staining Buffer Set (ThermoFisher Scientific) when intracellular staining was required. The following fluorochrome-conjugated antibodies were used for staining intracellular markers: FoxP3 (FJK-16s, eBioscience), granzyme B (NGZB, eBioscience), IFNγ (XMG1.2, Biolegend), Ki67 (SolA15, eBioscience) and TNFα (MP6-XT22, Biolegend). For flow cytometry analysis of myeloid cells, cells were stained with a combination of the following fluorochrome-conjugated antibodies: CD3ε (clone: 17A2), CD19 (clone: 6D5), TCRβ (clone: H57-597), NK1.1 (clone: PK136), F4/80 (clone: BM8), Ly6c (clone: HK1.4), Gr-1 (RB6-8C5), CD11b (M1/70), CD11c (clone: N418), IA/E (clone: M5/114.15.2), CD103 (clone: 2E7), CD8α (53-6.7), CD317 (clone: BST2, eBioscience), PDL-1 (clone: 10F.9G2), XCR1 (clone: S15046E), CD24 (clone: M1/69, BD Biosciences) and CD301b (clone: URA-1). Stained or fixed cells were resuspended in 300µl of FACS buffer and analyzed using a LSRFortessa X-20 flow cytometer. Data were analyzed using FlowJo software version 10.8.0.

### Mitochondrial metabolomics

sgNT and sg*Ndufs4* B16/BL6 cells were grown in four p-150 dishes to 80% confluent before being trypsinized and washed three times with cold PBS. The cell pellets were suspended in mitochondrial isolation buffer (0.32 M sucrose, 0.001 M EDTA and 0.01 M Tris with pH 7.2). The cell suspense was transferred into a dounce homogenizer and dounced 20 times and then subjected to centrifuge at 1000 g for 5 min at 4LJ°C. The supernatant was transferred to a new tube and centrifuged at 1000 g for 5 min at 4LJ°C again. The new obtained supernatant was the centrifuged at 9000 g for 15 min at 4LJ°C. The pellet was resuspended into mitochondrial isolation buffer and for protein concentration test with DC protein assay (Bio-Rad, 5000111). The obtained mitochondria were aliquoted to 1.5 ml Eppendorf tube with 100 ug per tube and centrifuged at 9000 g for 10 min. The obtained pellets were resuspended in 80% cold methanol and incubated in -80°C for one hour and were then centrifuged at maximum speed for 10 min at 4LJ°C. The top layer, which contains the polar metabolites, was removed, and transferred to a new tube and dried down overnight using a SpeedVac (Thermo Fisher Scientific). Lyophilized samples were resuspended in 20 μl HPLC quality water and subjected to metabolomics profiling using the AB/SCIEX 5500 QTRAP triple quadrupole instrument. Data analysis was performed using the GiTools software.

### Protein digestion and isobaric labelling for mass spectrometry analysis

Tumor samples from allograft study were lysed with 4 mL of SDS lysis buffer (2.0% SDS (w/v), 250 mM NaCl, 5 mM TCEP, EDTA-free protease inhibitor cocktail (Promega), and 100 mM HEPES, pH 8.5) using an Omni tissue homogenizer. Extracts were reduced at 57°C for 30 min and cysteine residues were alkylated with iodoacetamide (14 mM) in the dark at room temperature for 45 min. Extracts were purified by methanol–chloroform precipitation and pellets were washed with ice-cold methanol. Pellets were resuspended in 2 mL of 8 M urea (containing 50 mM HEPES, pH 8.5) and protein concentrations were measured by BCA assay (Thermo Fisher Scientific) before protease digestion. 100 µg of protein were diluted to 4 M urea and digested overnight with 4 µg LysC (Wako). Digests were diluted further to a 1.5 M urea concentration and 5 µg of trypsin (Promega) was added for 6 hours at 37°C. Digests were acidified with 50 µl of 20% formic acid and subsequently desalted by C18 solid-phase extraction (50 mg, Sep-Pak, Waters). Digested brain peptides were resuspended in 100 µl of 200 mM HEPES, pH 8.0. 10 μL of TMTpro reagents (Thermo Fisher Scientific) was added to each solution for 1 hour at room temperature (25°C). After incubating, the reaction was quenched by adding 4 μL of 5% (w/v) hydroxylamine. Labelled peptides were combined and subsequently desalted by C18 solid-phase extraction (50 mg, Sep-Pak, Waters) before basic pH reversed-phase separation.

### Basic pH reverse-phase separation for mass spectrometry analysis

Tandem mass tag (TMT)-labeled peptides were solubilized in 500 μL solution containing 5% acetonitrile in 10 mM ammonium bicarbonate, pH 8.0 and separated by an Agilent 300 Extend C18 column (5 mm particles, 4.6 mm inner diameter and 220 mm in length). An Agilent 1100 binary pump coupled with a photodiode array detector (Thermo Fisher Scientific) was used to separate the peptides. A 40-min linear gradient from 20% to 40% acetonitrile in 10 mM ammonium bicarbonate, pH 8 (flow rate of 0.6 ml/min) separated the peptide mixtures into a total of 96 fractions (30 sec). A total of 96 fractions was consolidated into 12 samples in a checkerboard fashion, acidified with 20 µl of 20% formic acid and vacuum dried to completion. Each sample was re-dissolved in 5% formic acid, 5% ACN, desalted via StageTips before liquid chromatograph–tandem mass spectrometry (LC–MS/MS) analysis. LC–MS/MS analysis. Data were collected using an Orbitrap Fusion Lumos mass spectrometer (Thermo Fisher Scientific, San Jose, CA) coupled with a Proxeon EASY-nLC 1200 LC pump (Thermo Fisher Scientific). Peptides were separated on a 100 μm inner diameter microcapillary column packed with 35 cm of Accucore C18 resin (2.6 μm, 100 Å, Thermo Fisher Scientific). Peptides were separated using a 3 hours gradient of 6–22% acetonitrile in 0.125% formic acid with a flow rate of ∼400 nL/min. Each analysis used an MS3-based TMT method as described previously described^44^. The data were acquired using a mass range of m/z 400 – 1400, resolution at 120,000, AGC target of 1 × 10^6^, a maximum injection time 100 ms, dynamic exclusion of 180 seconds for the peptide measurements in the Orbitrap. Data dependent MS 2 spectra were acquired in the ion trap with a normalized collision energy (NCE) set at 35%, AGC target set to 2.0 × 10^5^ and a maximum injection time of 120 ms. MS3 scans were acquired in the Orbitrap with an HCD collision energy set to 45%, AGC target set to 1.5 × 10^5^, maximum injection time of 200 ms, resolution at 50,000 and with a maximum synchronous precursor selection (SPS) precursors set to 10.

### Mass spectrometry data processing and spectra assignment

In-house developed software was used to convert acquired mass spectrometric data from the .RAW file to the mzXML format. Erroneous assignments of peptide ion charge state and monoisotopic m/z were also corrected by the internal software. SEQUEST algorithm was used to assign MS2 spectra by searching the data against a protein sequence database including Mouse Uniprot Database (downloaded June 2018) and known contaminants such as mouse albumin and human keratins. A forward (target) database component was followed by a decoy component including all listed protein sequences. Searches were performed using a 20ppm precursor ion tolerance and requiring both peptide termini to be consistent with trypsin specificity. 16-plex TMT labels on lysine residues and peptide N termini (+304.2071 Da) were set as static modifications and oxidation of methionine residues (+15.99492 Da) as a variable modification. An MS2 spectra assignment false discovery rate (FDR) of less than 1% was implemented by applying the target-decoy database search strategy. Filtering was performed using a linear discrimination analysis method to create one combined filter parameter from the following peptide ion and MS2 spectra properties: XCorr and ΔCn, peptide ion mass accuracy, and peptide length. Linear discrimination scores were used to assign probabilities to each MS2 spectrum for being assigned correctly and these probabilities were further used to filter the data set with an MS2 spectra assignment FDR to obtain a protein identification FDR of less than 1%.

### Determination of TMT reporter ion intensities

For reporter ion quantification, a 0.003 m/z window centered on the theoretical m/z value of each reporter ion was monitored for ions, and the maximum intensity of the signal to the theoretical m/z value was recorded. Reporter ion intensities were normalized by multiplication with the ion accumulation time for each MS2 or MS3 spectrum and adjusted based on the overlap of isotopic envelopes of all reporter ions. Following extraction of the reporter ion signal, the isotopic impurities of the TMT reagent were corrected using the values specified by the manufacturer’s specification. Total signal-to-noise values for all peptides were summed for each TMT channel and all values were adjusted to account for variance and a total minimum signal-to-noise value of 200 was implemented.

### Statistical analysis and reproducibility

All measurements were biological replicate samples. In general, for two experimental comparisons, unpaired two-tailed Mann-Whitney u-test was used. GraphPad Prism software (version 10.1.0) and R were used to generate graphs and perform statistical analyses. Microsoft Excel was used for analysis of small-molecule screen data and proteomics. DAVID Bioinformatics Resources 6.8 was used for gene ontology analysis^45,46^. TCGA survival data was analysized using the https://www.tcga-survival.com/ tool^47^. Immunotherapy response datasets were accessed and analyzed through TIGER^48^.

---

**Extended Data Fig. 1:**
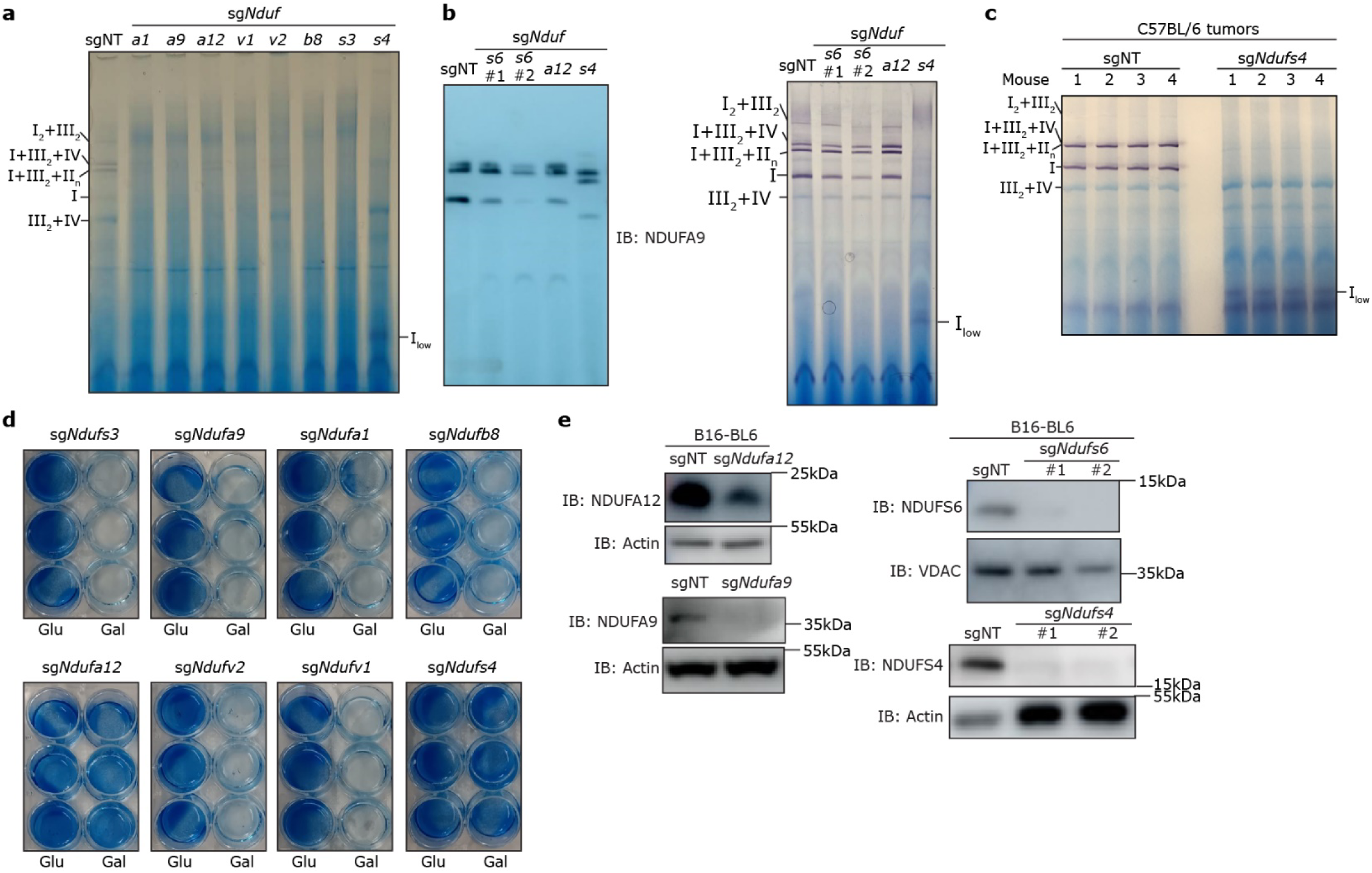
CI in gel activity assay and galactose sensitivity test for CI subunit knockouts. **a**, Blue native page gel showing CI activity from isolated crude mitochondria of CI subunit CRISPR knockouts. Purple bands indicate CI activity, blue bands indicate protein presence from Coomassie. Supercomplexes are labeled based on previous work^51^. **b**, Blue native page gel showing CI activity in sg*Ndufs6*, sg*Ndufa12*, and sg*Ndufs4* cells and western blot showing NDUFA9 assembly into CI supercomplexes in sg*Ndufa12* and sg*Ndufs4* cells. Different numbers under sg*Ndufs6* represent different CRISPR guides. **c**, Blue native page gels showing CI activity from isolated crude mitochondria from sgNT and sg*Ndufs4* tumors from C57BL/6 mice. **d**, Cell titer blue of CI knockout cells grown in either glucose (Glu) or galactose (Gal). **e**, Western blots showing knockouts for sg*Ndufa12,* sg*Ndufa9*, sg*Ndufs6* and sg*Ndufs4* cell lines.

**Extended Data Fig. 2:**
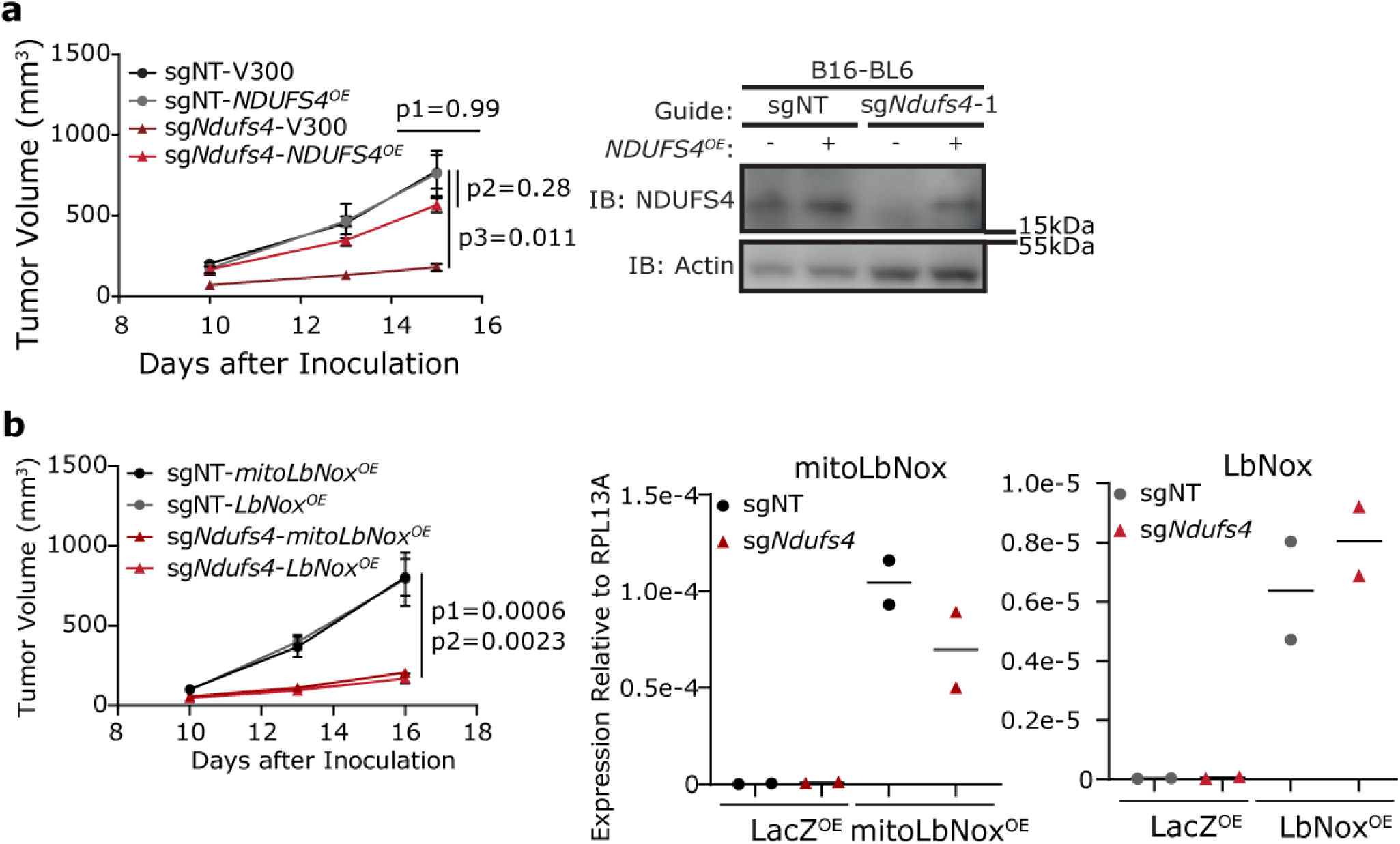
Anti-tumor effects of sg*Ndufs4* cells are rescued by *Ndufs4* overexpression and are not dependent on the redox state of the cell. **a**, Growth curves of sgNT and sg*Ndufs4* tumors transduced with either an empty or a *Ndufs4* over expression vector in C57BL/6 mice. P1 compares sgNT-V300 to sgNT-*Ndufs4^OE^*, P2 compares sgNT-*Ndufs4^OE^*to sg*Ndfus4-Ndufs4^OE^*, P3 compares sgNT-V300 to sg*Ndufs4*-V300. Western blot confirmation of *Ndufs4* over expression. **b**, Growth curves of sgNT and sg*Ndufs4* tumors transduced with either an *LbNox* or mitochondrial target *LbNox* over expression vector in C57BL/6 mice. P1 compares sgNT-*mitoLbNox^OE^* to sg*Ndufs4*-*mitoLbNox^OE^*and p2 compares sgNT-LbNox^OE^ to sg*Ndufs4*-*LbNox^OE^*. qPCR validation of *LbNox* overexpression. N = 5 mice per condition in **a** and **b**. *P* values in **a** and **b** were calculated using Brown-Forsythe and Welch ANOVA test with Dunnett’s T3 multiple comparisons test. Error bars are mean□±□s.e.m.

**Extended Data Fig. 3:**
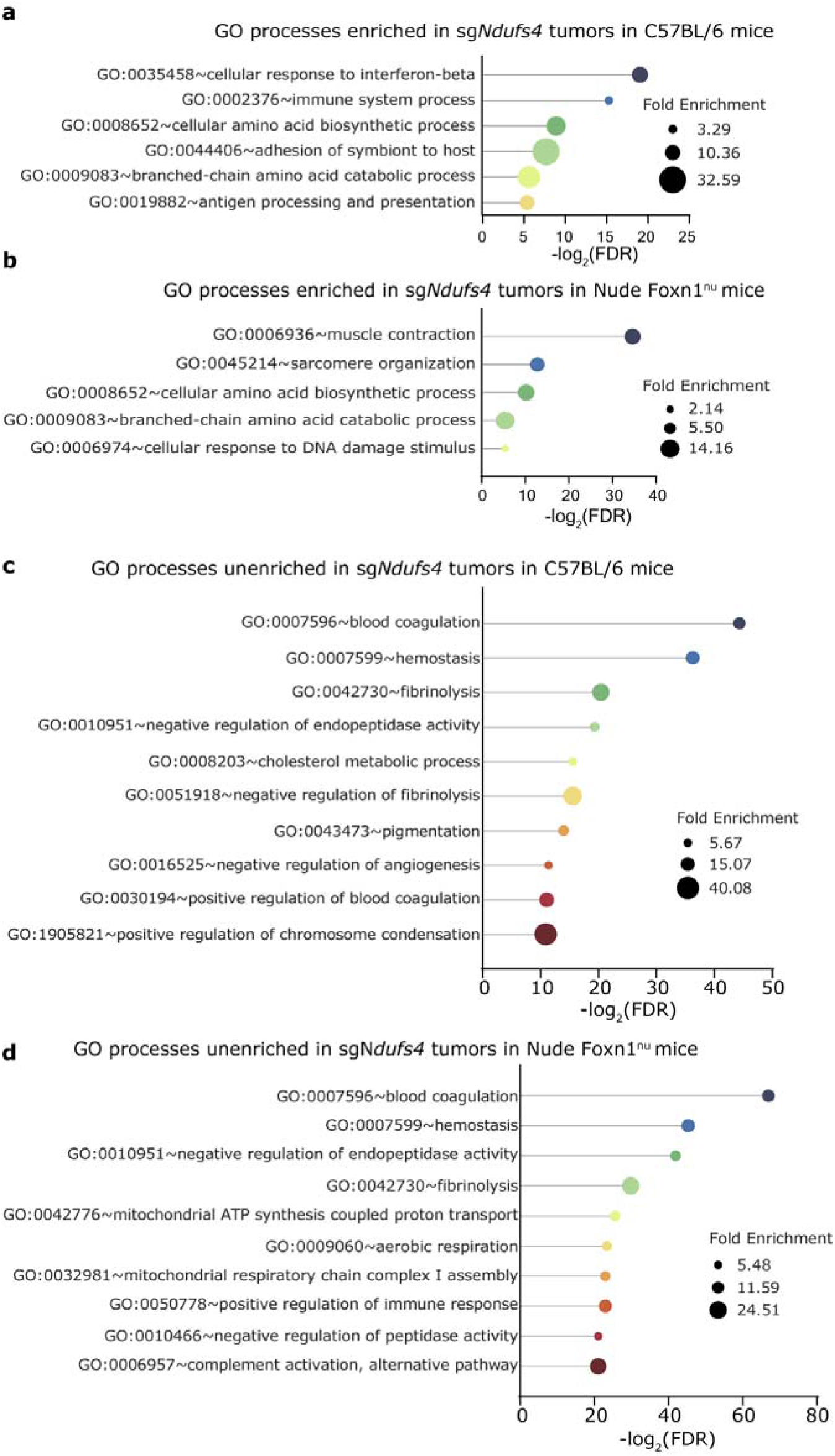
Gene Ontology analysis of tumor proteomics from C57BL/6 and nude mice using DAVID^45,46^. **a**, Enriched gene sets in sg*Ndufs4* tumors in C57BL/6 mice using proteins with fold-change ≥1.5. **b**, Enriched gene sets in sg*Ndufs4* tumors in nude mice using proteins with fold-change ≥1.5. **c**, Unenriched gene sets in sg*Ndufs4* tumors in C57BL/6 mice using proteins with fold-change ≥1.5. **d**, Unenriched gene sets in sg*Ndufs4* tumors in nude mice using proteins with fold-change ≥1.5.

**Extended Data Fig. 4:**
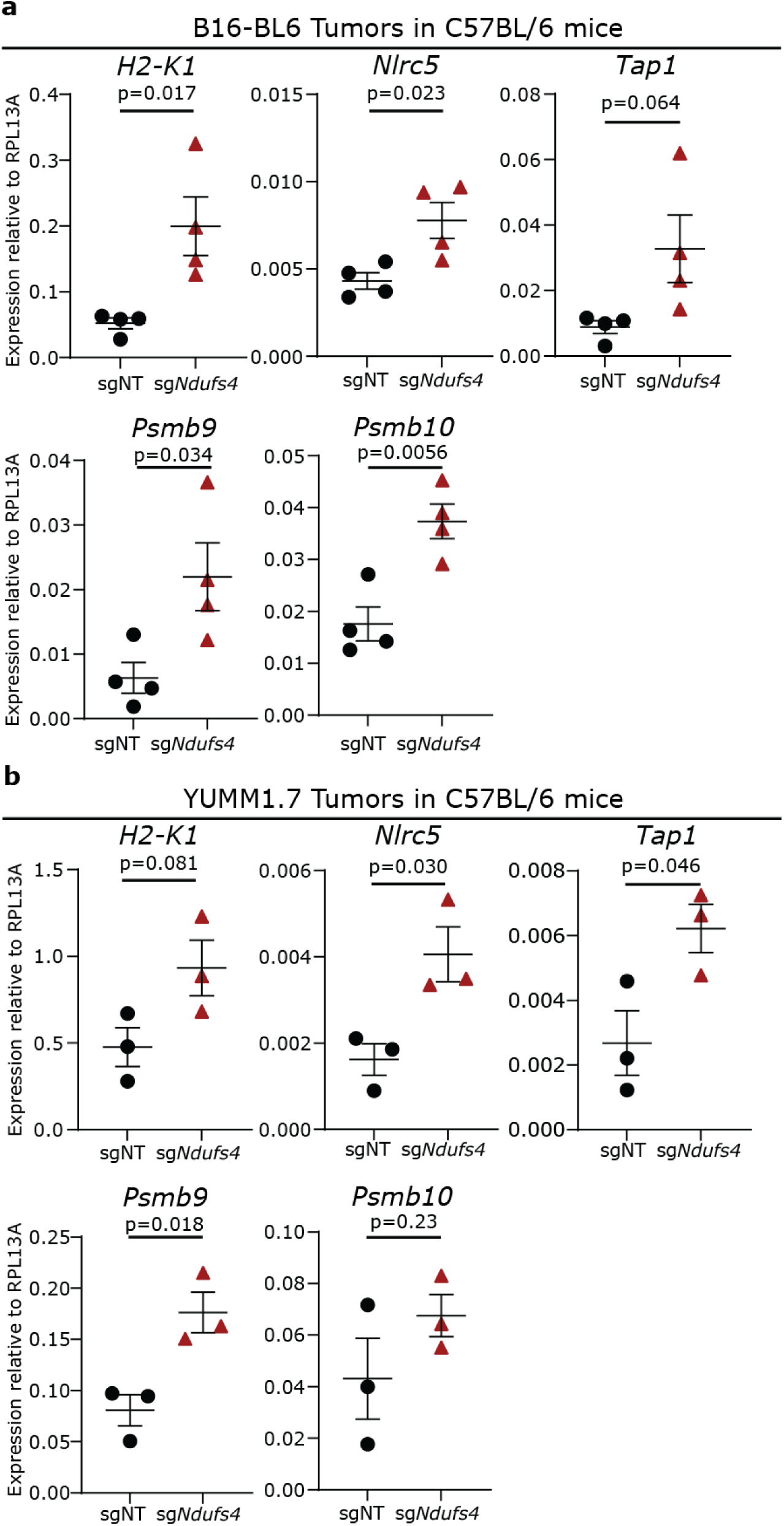
mRNA levels of antigen processing and immune related genes from tumor grown in C57BL/6 mice. **a**, Expression levels of MHC-I and antigen processing genes from sgNT or sg*Ndufs4* B16-BL6 tumors from C57BL/6 mice (n=4 per condition). **b**, Expression levels of MHC-I and antigen processing genes from sgNT or sg*Ndufs4* YUMM1.7 tumors form C57BL/6 mice (n=3 per condition). *P* values in **a** and **b** were calculated using a two-tailed, unpaired Student’s t-test. Error bars are mean□±□s.e.m.

**Extended Data Fig. 5:**
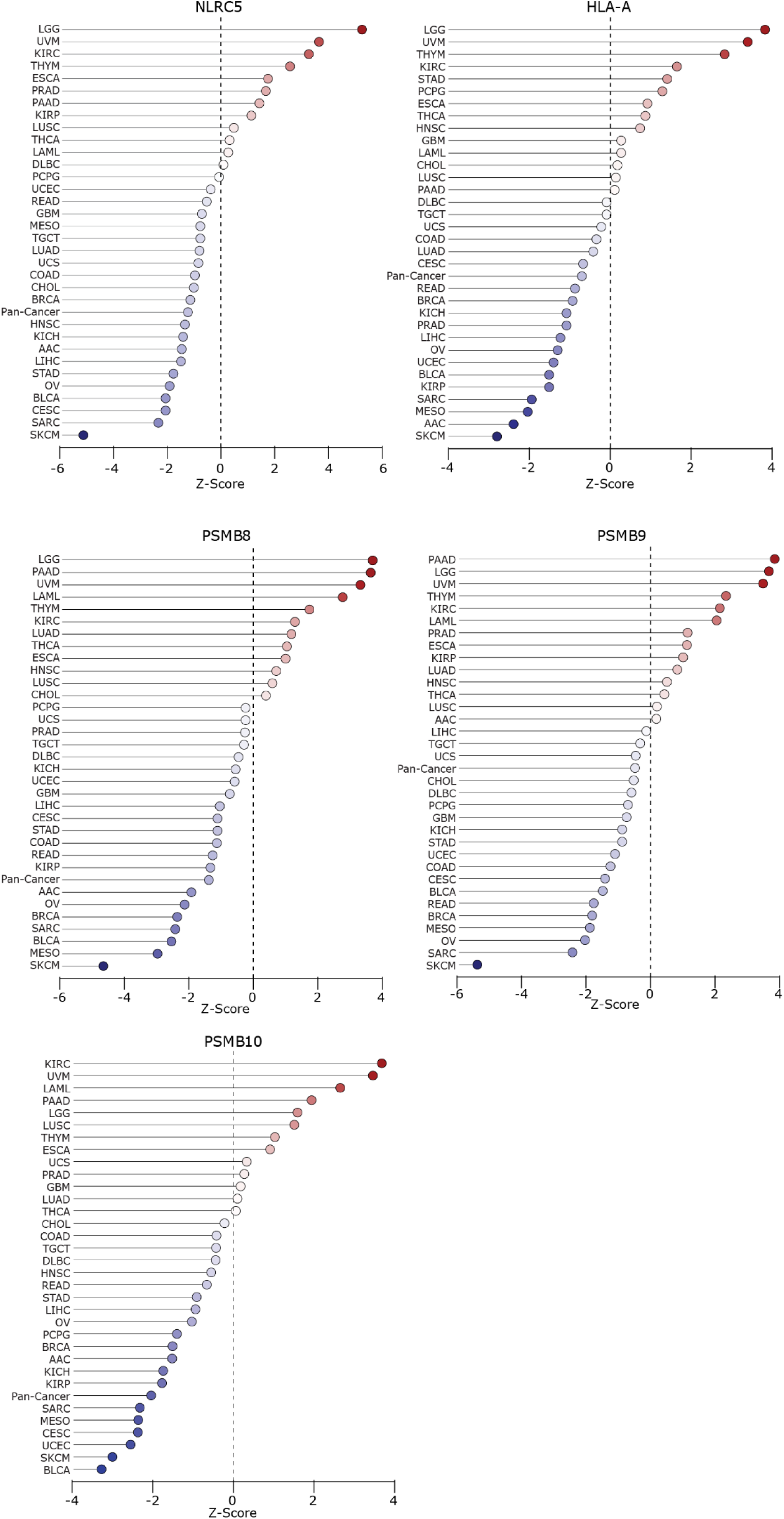
sg*Ndufs4* antigen processing and presentation signature correlates most with survival in SKCM dataset. TCGA Z-score survival analysis ranking of all TCGA cancer types for antigen processing and presentation genes found to be upregulated in sg*Ndufs4* tumors (*NLRC5, HLA-A, PSMB8, PSMB9*, and *PSMB10*). Low z-scores indicate that high expression of a gene is associated with better prognosis, high z-scores indicate that high expression of a gene is associated with worse prognosis.

**Extended Data Fig. 6:**
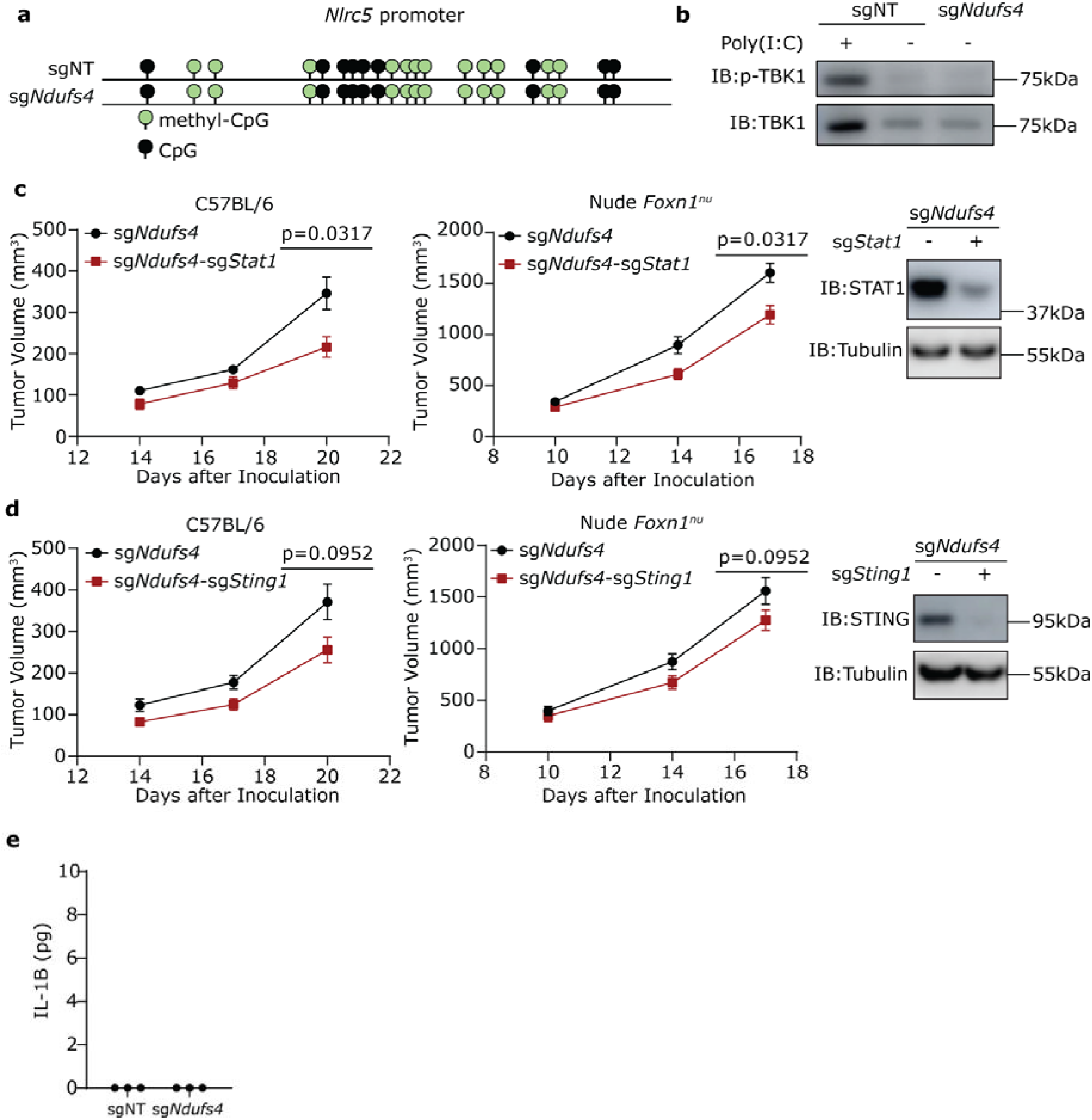
Nlrc5 gene regulation isn’t associated with increased promoter methylation or regulation of NF-κB downstream of cGAS-STING or STAT1. **a**, CpG viewer output of bisulfate sequencing of the *Nlrc5* promoter region in sgNT and sg*Ndufs4* cells in vitro. **b**, Western blot analysis of TBK1 and phospho-TBK1 protein levels in sgNT and sg*Ndufs4* cells in vitro. **c**, Growth curves of sg*Ndufs4* and sg*Ndufs4*-sg*Stat1* tumors along with western blot of protein levels. **d**, Growth curves of sg*Ndufs4* and sg*Ndufs4*-sg*Sting1* tumors along with western blot of protein levels. **e**, IL-1B concentration in cell culture media from sgNT and sgNdufs4 cells. N=5 mice per condition for **b-c**. *P* values in **b**-**c** were calculated using a Mann Whitney *u*-test. Error bars are mean□±□s.e.m.

**Extended Data Fig. 7:**
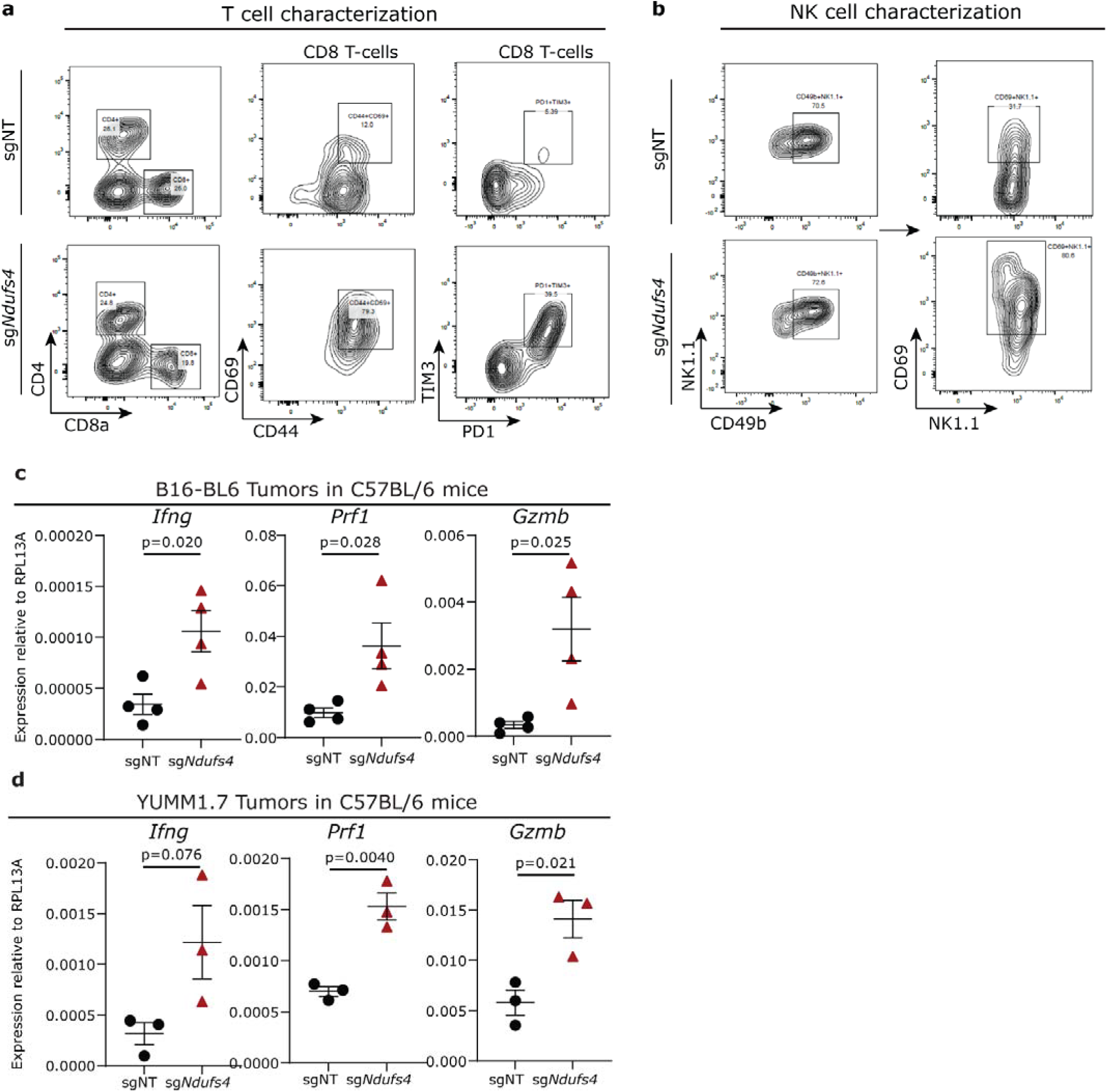
Gating strategy for T-cell and NK cells in sgNT and sg*Ndufs4* tumors. **a**, Density plots showing CD8 versus CD4 discrimination, expression of activation markers CD44 and CD69 and expression of exhaustion markers PD-1 and TIM-3 on CD8 T-cells. **b**, Density plots showing NK cell discrimination using markers CD49b and NK1.1 and expression of the activation marker CD69 on NK cells. Plots are representative examples from experiments. **c**, Expression of immune related genes from B16-BL6 tumors grown in C57BL/6 mice (n=4 mice per condition). **d**, Expression of immune related genes from YUMM1.7 tumors grown in C57BL/6 mice (n=3 mice per condition).

